# Improved transformation efficiency of group A *Streptococcus* by inactivation of a type I restriction modification system

**DOI:** 10.1101/2021.02.23.432441

**Authors:** Meredith B. Finn, Kathryn M. Ramsey, Simon L. Dove, Michael R. Wessels

**Affiliations:** Division of Infectious Diseases, Boston Children’s Hospital and Department of Pediatrics, Harvard Medical School, Boston, MA, USA; Departments of Cell and Molecular Biology and Biomedical and Pharmaceutical Sciences, University of Rhode Island, Kingston, Rhode Island, United States of America

## Abstract

*Streptococcus pyogenes* or group A *Streptococcus* (GAS) is a leading cause of bacterial pharyngitis, skin and soft tissue infections, life-threatening invasive infections, and the post-infectious autoimmune syndromes of acute rheumatic fever and post-streptococcal glomerulonephritis. Genetic manipulation of this important pathogen is complicated by resistance of the organism to genetic transformation. Very low transformation efficiency is attributed to recognition and degradation of introduced foreign DNA by a type I restriction-modification system encoded by the *hsdRSM* locus. DNA sequence analysis of this locus in ten GAS strains that had been previously transformed with an unrelated plasmid revealed that six of the ten harbored a spontaneous mutation in *hsdR, S*, or *M*. The mutations were all different, and at least five of the six were predicted to result in loss of function of the respective *hsd* gene product. The unexpected occurrence of such mutations in previously transformed isolates suggested that the process of transformation selects for spontaneous inactivating mutations in the Hsd system. We investigated the possibility of exploiting the increased transformability of *hsd* mutants by constructing a deletion mutation in *hsdM* in GAS strain 854, a clinical isolate representative of the globally dominant M1T1 clonal group. Mutant strain 854Δ*hsdM* exhibited a 5-fold increase in transformation efficiency compared to the wild type parent strain and no obvious change in growth or off-target gene expression. We conclude that genetic transformation of GAS selects for spontaneous mutants the *hsdRSM* restriction modification system. We propose that use of a defined *hsdM* mutant as a parent strain for genetic manipulation of GAS will enhance transformation efficiency and reduce the likelihood of selecting spontaneous *hsd* mutants with uncharacterized genotypes.

## Introduction

As part of our research into molecular pathogenesis of infections due to group A *Streptococcus* (*S. pyogenes* or GAS), our laboratory routinely constructs mutant strains of GAS using allelic exchange mutagenesis by transformation of a wild type strain with a temperature-sensitive plasmid carrying the desired mutation. During characterization of one such mutant strain, we made the incidental discovery that the mutant harbored a 694 bp deletion in *hsdM*, which encodes a component of a type I restriction modification system. The deletion resulted in a frameshift mutation at codon 123, altering the remainder of the protein sequence and introducing a premature stop codon at amino acid 170. As discussed below, because inactivation of this system can be associated with enhanced susceptibility to transformation with foreign DNA, three questions arose concerning inactivating mutations in *hsdM* or in genes encoding other components of the restriction modification system: (1) Does inactivation of *hsdM* result in increased transformation efficiency? (2) Does the process of transformation select for spontaneous inactivating mutations in the restriction modification system? (3) Could deliberate disruption of the system be exploited to facilitate introduction of foreign DNA for genetic manipulation of GAS? The current investigation was undertaken to answer these questions.

Type I restriction modification systems are comprised of three subunits that work together to recognize and cleave intracellular foreign DNA at specific sites and also to protect host cell DNA from cleavage by methylating it at the same recognition sites. Together, the three protein subunits comprise the “<u>h</u>ost <u>s</u>pecificity of <u>D</u>NA” system, or Hsd. The three enzymes of the Hsd system include the specificity subunit, HsdS, the restriction subunit, HsdR, and the modification subunit, HsdM. HsdS requires interaction with HsdM in order to bind to DNA. Once bound, the HsdS/HsdM complex can interact with HsdR. HsdR can then exert endonuclease activity at specific recognition sites, activity which is dependent on the methylation state of the target sequence. These functions of the three enzymes are tightly linked, as interaction with HsdM is necessary both for HsdS to bind to DNA and for HsdR endonuclease activity to occur [1]. Together, the three-protein complex is able to cleave foreign DNA after it enters the cell as a type of bacterial defense system.

The Hsd system in GAS is a typical type I restriction modification system. It efficiently targets and cleaves foreign DNA that may enter the cell either naturally or artificially through electroporation. Accordingly, GAS is notoriously difficult to manipulate genetically in the laboratory setting. Previous studies have reported that inactivation of the Hsd system in GAS is associated with increased transformation efficiency [2,3]. One study done in an M28 strain found that deletion of the three components of the Hsd system was associated with increased transformation efficiency [2]. Another study characterized a group of *emm1* clinical isolates associated with invasive infections. These strains were found to harbor a spontaneous deletion that included part of *hsdR*, the entire *hsdS* and *hsdM* gene sequences, and a portion of an adjacent two-component system. Strains with the deletion were shown to have increased transformation efficiency compared to contemporaneous *emm1* isolates without the deletion [3].

In this study, we introduced a large deletion in *hsdM* in a GAS strain representative of the M1T1 clonal group widely implicated in invasive GAS disease. We found that inactivation of *hsdM* resulted in increased transformation efficiency of the strain, and that the mutation was not associated with a significant change in transcript abundance for other GAS genes including those encoding key virulence factors. In addition, we observed that the process of transformation selects for strains that have acquired spontaneous mutations in the Hsd system. We suggest that using a defined *hsdM* mutant as the parental strain for genetic manipulations could improve transformation efficiency and reduce the likelihood that transformation would select for an undefined spontaneous mutation in the restriction modification system with unknown potential off-target effects.

## Results

### Inactivation of *hsdM* results in increased transformation efficiency

Since restriction modification systems target foreign DNA for destruction by the host cell, we hypothesized that inactivation of the Hsd system by deletion of *hsdM* would result in a strain with increased transformation efficiency. To test this hypothesis, we introduced a 1,005 bp in-frame deletion into the *hsdM* coding sequence of GAS strain 854, an *emm1* isolate representative of the globally prevalent M1T1 clonal group that is dominant among GAS *emm1* strains associated with invasive infections [4]. The mutation of *hsdM* was confirmed by DNA sequencing. No growth deficiency was observed in a growth curve compared to wild type strain 854 grown in THY medium. We tested transformation efficiency by electroporation of wild type strain 854 or the *hsdM* knockout strain 854Δ*hsdM* using 1 µg of plasmid DNA. We then serially diluted the transformation reactions, plated the dilutions on selective agar, and counted the number of transformants that resulted from each reaction. These experiments revealed an increase in transformation efficiency of the Δ*hsdM* strain of approximately 5-fold compared to the wild type strain (Figure 1).

**Figure 1.**
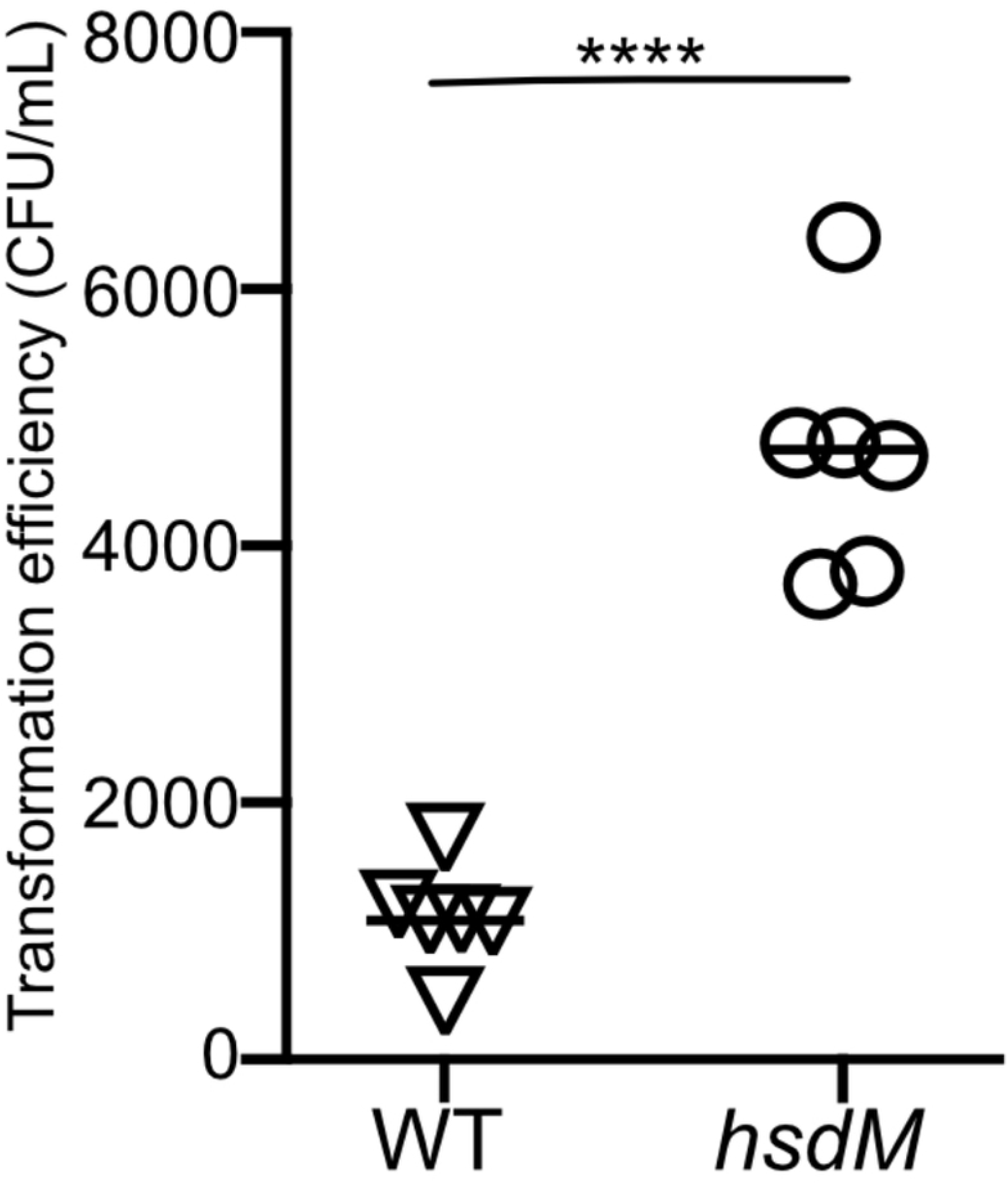
Deletion of *hsdM* results in increased transformation efficiency. Cells of wild type GAS strain 854 (triangles) or mutant strain 854Δ*hsdM* (circles) were subjected to electroporation with 1 µg of plasmid DNA and then serially diluted and spread on selective medium for quantitative culture. Data points represent independent experiments; horizontal bars indicate the mean. \*\*\*\**P*<0.0001, Student’s t-test.

### Transformation of wild type GAS selects for mutants in *hsdRSM*

Because inactivation of HsdM resulted in a higher transformation rate, we wondered whether the process of transformation of wild type GAS might select for bacterial cells that had acquired a spontaneous inactivating mutation in the Hsd system. We screened for such mutations by sequence analysis of the *hsdM* locus of ten independently derived mutant strains of GAS 854 that had been previously transformed during allelic exchange mutagenesis of various genes. We found that four of the ten had acquired spontaneous mutations in the *hsdM* gene. These mutations included a deletion encompassing the entire *hsdM* coding sequence, a 1.5kb insertion, a missense point mutation, and a nonsense mutation (Table 1). Sequence analysis of the entire *hsdRSM* locus revealed that two other strains among these ten had mutations in either *hsdR* or *hsdS* (Table 1). Thus, including the mutations in *hsdM*, we found spontaneous mutations in the *hsdRSM* locus in six of the ten strains that had been previously transformed. Five of the six mutations introduced premature stop codons and/or large insertions or deletions in coding regions, strongly suggesting that these mutations all result in inactivation of at least one component of the Hsd system. By modeling the structure of HsdM to the published structure of a homolog in *E. coli* (PDB accession number 7BST) [5], we mapped amino acid T316 to the interface between HsdM and HsdR (Figure 2). The corresponding residue in the *E*. coli HsdM, N319, has been proposed to interact with R456 of HsdR, which aligns with K449 in the GAS HsdR sequence. Thus, mutation of GAS T316 from threonine to proline in the CsrS D148N E151Q D152N strain may disrupt the interaction of HsdM with HsdR and impact the function of the enzyme complex.

**Table 1.** Spontaneous mutations in *hsdRSM* found in isolates of GAS strain 854 that have previously been subjected to genetic transformation.

|  |  |
| --- | --- |
| **Strain** | ***hsd* mutation** |
| 854Δ*csrR* | 694 bp deletion in the coding region of *hsdM* |
| 854CsrS H280A | Premature stop codon at position 380 (of 526) in *hsdM* |
| 854Slo Y255A | 1.5 kb insertion in the coding region of *hsdM* |
| 854CsrS D148N<br>E151Q D152N | T316P substitution in HsdM |
| 854CsrS T284A | 4 nucleotide insertion duplication that introduces a premature stop codon at position 794 (of 992) in *hsdR* |
| 854NADase G330D | 621 bp deletion in the coding region of *hsdS* resulting in a deletion from E160 to G367 |
| 854Δ*slo* | none |
| 854Δ*nga* | none |
| 854Δ*csrS* | none |
| 854*speB::*$\Omega$ | none |

**Figure 2.**
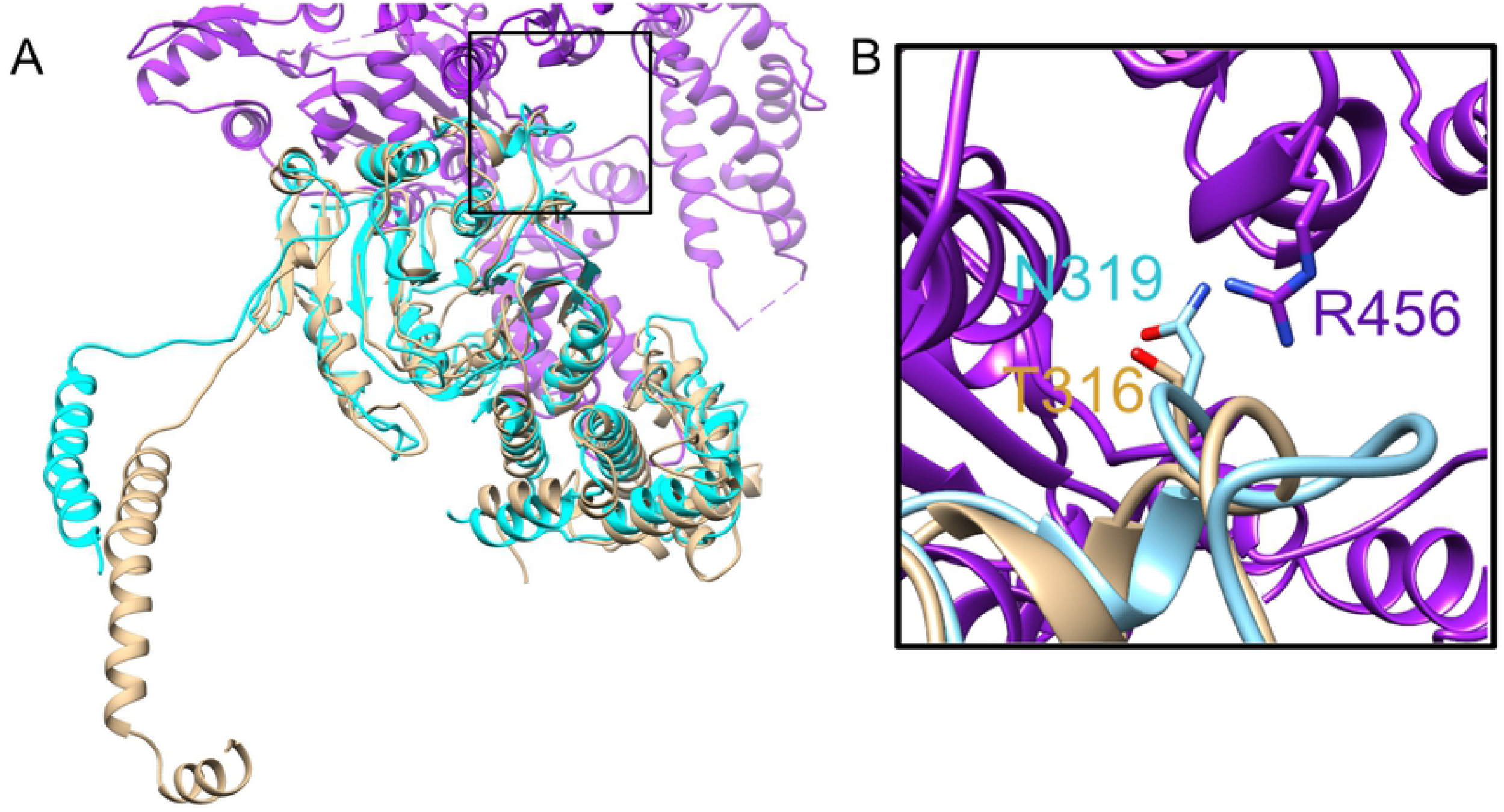
Modeling of GAS HsdM. **(A)** Modeled structure of HsdM from GAS is shown in brown superimposed on a portion of the published structure of HsdR_2_M_2_S_1_-Ocr^IS^ from EcoR124I (PDB 7BST). *E. coli* HsdM is shown in teal, and HsdR is shown in purple. **(B)** Close up view of boxed section of panel A showing the interface between HsdM and HsdR. Proteins are in ribbon representation with residues R456 from HsdR, N319 from HsdM (EcoR124I) and the corresponding residue T316 from HsdM (GAS) shown in stick representation. Alignment has a normalized Z-score of 3.30.

Since each of the spontaneous mutations we observed in transformants of strain 854 is different, we infer that the various mutations in the *hsdRSM* locus arose as independent events, rather than a single ancestral mutation in the parental wild type strain. Taken together, these observations strongly suggest that GAS isolates that have been successfully transformed in the laboratory setting may have a mutation that inactivates the Hsd system, and that the process of transformation selects for such mutants.

### Deletion of *hsdM* in *emm1* GAS strain 854 is not associated with altered expression of virulence genes

To assess the impact of inactivating *hsdM* in strain 854, we performed RNA-seq to evaluate global gene expression changes in the Δ*hsdM* strain compared to wild type. Twelve genes exhibited a 2-fold or greater change in transcript abundance for the mutant compared to wild type (Table 2). Assessment of gene expression by qRT-PCR of these twelve genes failed to confirm a significant change in expression of any of the genes (Table 2).

**Table 2.**
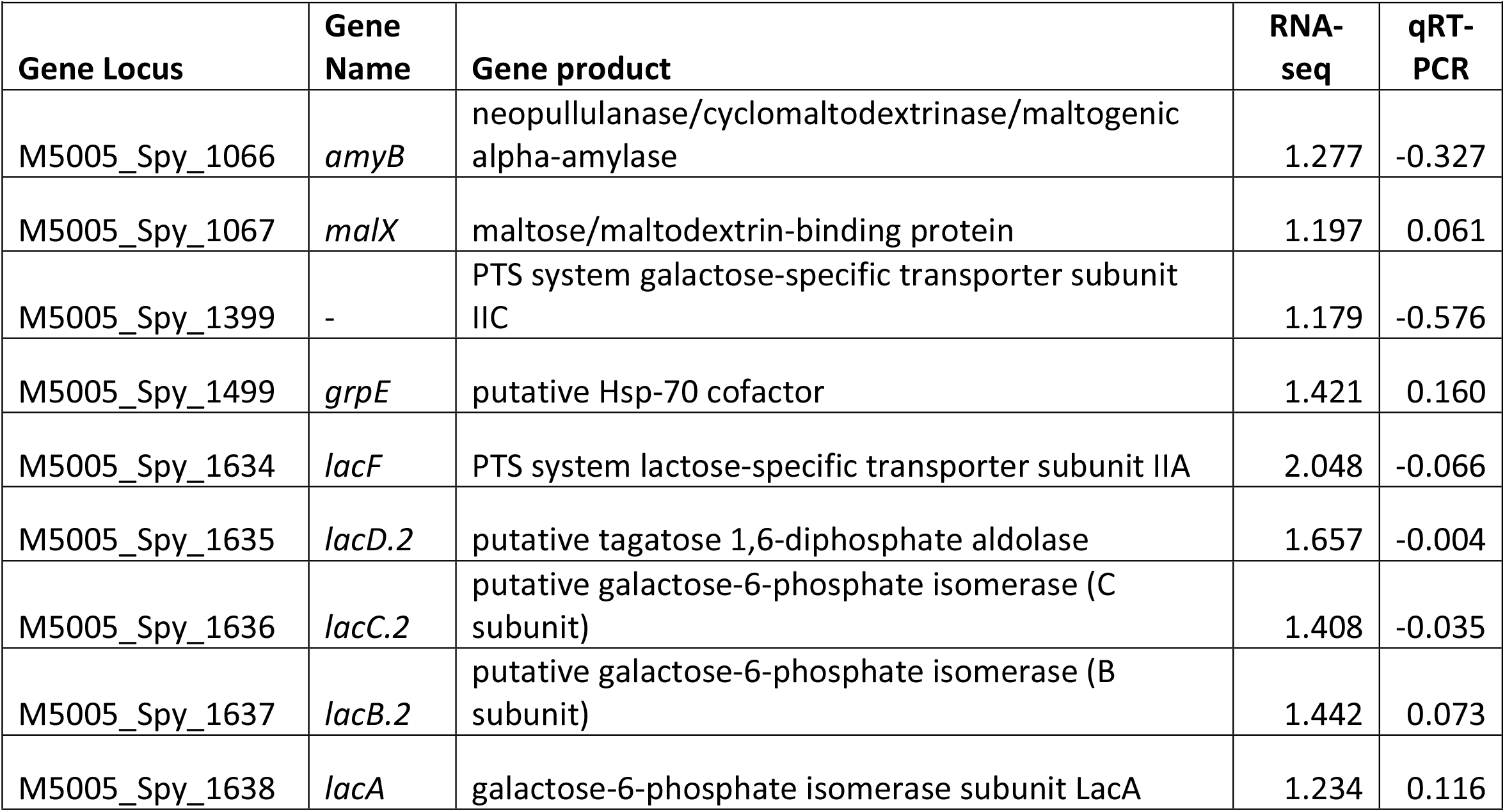
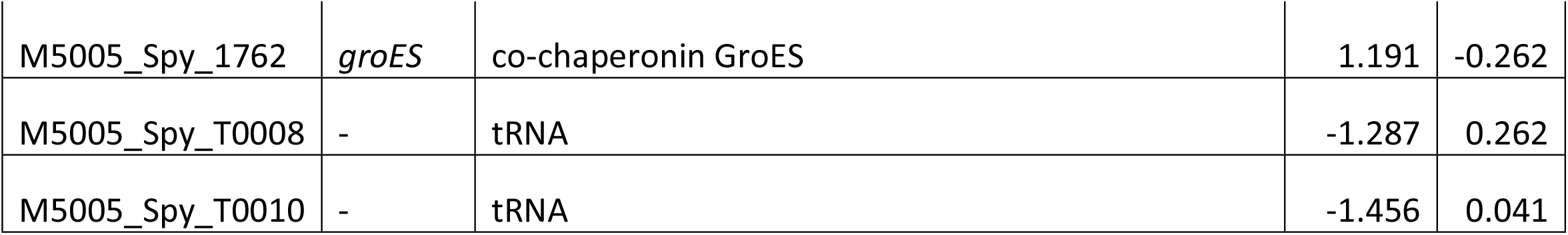
RNA-seq analysis of strain 854Δ*hsdM*.

**Table 2.** Changes in gene expression in strain 854Δ*hsdM* as assessed by RNA-seq or qRT-PCR. Results are shown as log_2_ fold-change in 854Δ*hsdM* relative to wild type strain 854.

A previous study reported that deletion of the *hsdRSM* locus in an M28 GAS strain resulted in reduced transcription of *mga*, which encodes a positive regulator of genes encoding several virulence factors including M protein (*emm28*) and C5a peptidase (*scpA*) [2]. To determine the impact of *hsdM* deletion on the transcript abundance of key virulence factors in the M1 strain 854, we used qRT-PCR to compare relative expression of genes encoding nine virulence factors in 854 wild type and 854Δ*hsdM* mutant cells. We observed no significant difference in the transcript abundance of these genes at either exponential or stationary growth phase (Figure 3), results that suggest inactivation of *hsdM* in strain 854 does not impact the regulation of major virulence factors including *mga* and Mga-regulated genes. Taken together, these results suggest that inactivation of *hsdM* in strain 854 does not significantly change expression of other genes in the 854 genome.

**Figure 3.**
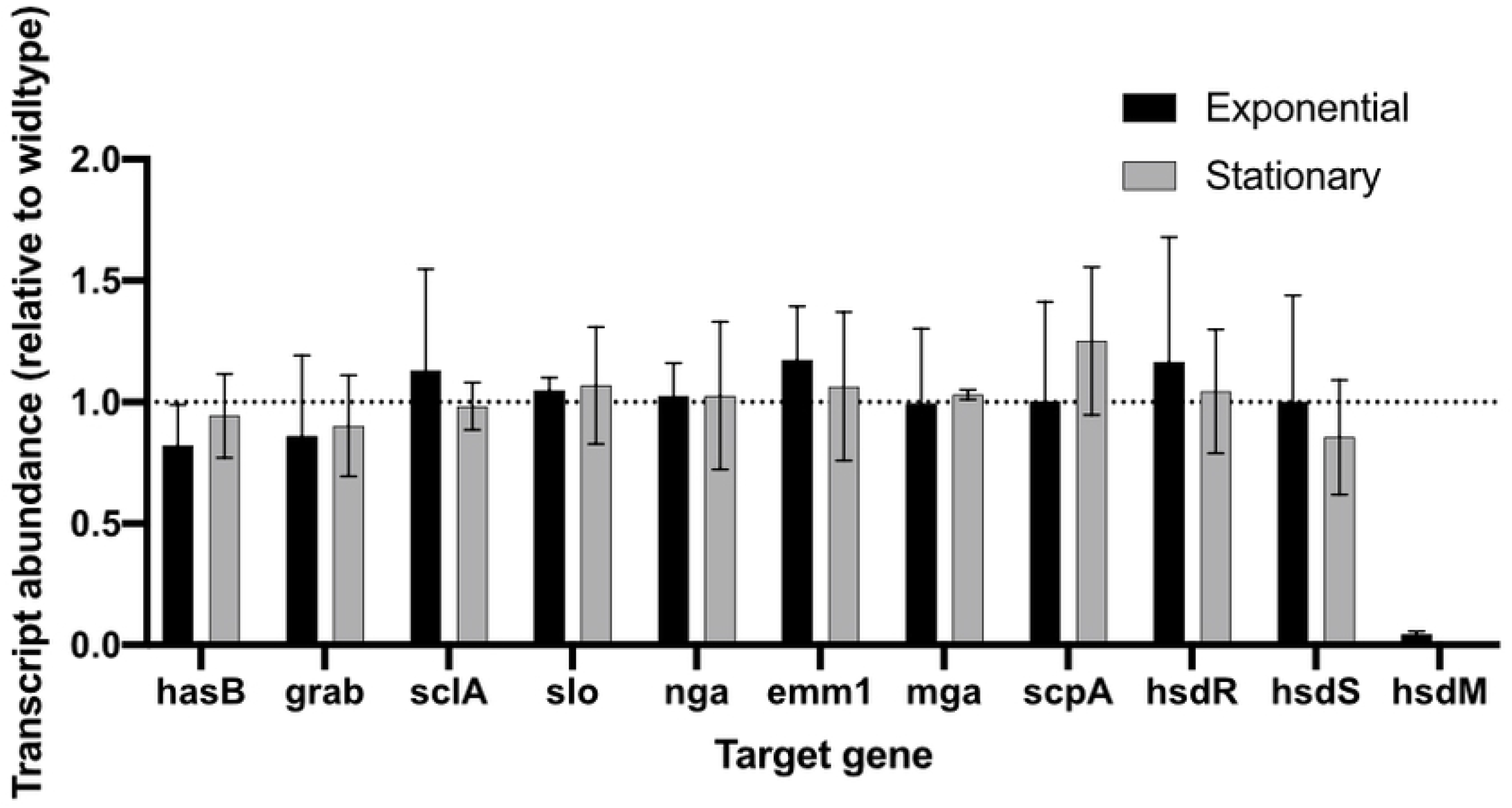
Transcriptional analysis of selected virulence factors in mutant strain 854Δ*hsdM*. Data represent mean ±SD transcript abundance as determined by qRT-PCR, expressed as a ratio of 854Δ*hsdM* to wild type strain 854. RNA was isolated from exponential (black) or stationary (gray) phase cultures.

## Discussion

In this study we report that the deletion of *hsdM* in the M1 GAS strain 854 results in increased transformation efficiency due to inactivation of the Hsd restriction modification system. We also observed that the process of transformation of a wild type strain selects for spontaneous mutants in *hsdRSM*, and that mutations in this locus appear to arise frequently and in a variety of ways. A global analysis of transcript levels in the *hsdM* deletion strain revealed no significant changes in expression of other genes compared to wild type. A previous study in an M28 strain found that deletion of the entire *hsdRSM* locus was associated with reduced expression of *mga* and Mga-regulated genes. That we did not detect changes in transcription of these or other genes may reflect strain- or serotype-specific differences or effects of deleting *hsdM* alone in contrast to the entire *hsd* locus.

Various approaches and modifications to restriction-modification systems have been utilized to increase transformation efficiencies in a variety of bacterial species. Inactivation of the Sau1 restriction modification system in *Staphylococcus aureus* strain RN4220 allows this strain to accept plasmid DNA from *E. coli* [6]. Plasmid DNA isolated from RN4220 can be further transformed among other *S. aureus* strains. Modification of a plasmid to eliminate the recognition sequences of restriction modification systems also has resulted in increased ability to transform *S. aureus* strains without having to use the RN4220 intermediate [7]. Deletion of the gene encoding the restriction endonuclease of a restriction modification system in *Caldicellulosiruptor* resulted in the ability to take up unmodified plasmid DNA from *E. coli* and transformation of plasmids isolated from different *Caldicellulosiruptor* species [8].

In light of the high frequency and variety of naturally occurring mutations we observed in the *hsdRSM* locus in strains that have undergone transformation in a laboratory setting, we believe it may be advantageous to use a defined Δ*hsdM* mutant as the parental GAS strain for further genetic manipulations. Use of a defined Δ*hsdM* mutant such as the one we describe here increases the likelihood of successful transformation, reduces the chance of selecting for spontaneous, uncharacterized *hsdRSM* mutants, and, in strain 854, appears not to result in significant changes in expression of other genes.

## Methods

### Bacterial strains and growth conditions

The GAS strain used for molecular manipulation was 854, an M1 strain isolated from a patient with a retroperitoneal abscess [9]. Mutants of strain 854 used in this study are listed in Table 3. GAS was cultured on trypticase soy agar supplemented with 5% defibrinated sheep blood (Remel) or Todd-Hewitt (TH) agar or in TH yeast (THY) broth at 37°C in the presence of 5% CO_2_. *E. coli* strain NEB5α (New England Biolabs) was used for cloning. *E. coli* was cultured in LB broth (Sigma). Growth media were supplemented with spectinomycin at 100 μg/ml when

**Table 3.** GAS strains used in this study.

| Table 3. GAS strains used in this study |  |  |
| --- | --- | --- |
| *S. pyogenes* strain | Description | Reference |
| 854 | Wild type M1 | [9] |
| 854$\Delta hsdM$ | 1,005 bp deletion in the coding sequence of *hsdM* | this study |
| 854$\Delta csrS$ | deletion of *csrS* | [10] |
| 854CsrS D148N E151Q D152N | Three point mutations in the predicted extracellular domain of CsrS | [11] |
| 854$\Delta csrR$ | Deletion of *csrR* | [11] |
| 854SLO Y255A | Y255A mutation in SLO | [12] |
| 854Δ*slo* | Deletion of *slo* | [10] |
| 854*speB::*$\Omega$ | Insertional inactivation of *speB* | [13] |
| 854CsrS H280A | H280A mutation in CsrS | [11] |
| 854CsrS T284A | T284A mutation in CsrS | [14] |
| 854Δ*nga* | Deletion of *nga* | [12] |
| 854*nga* G330D | G330D mutation in NADase | [12] |

### Construction of allelic exchange vector and GAS mutagenesis

Oligonucleotide primers are listed in Table S1. To generate the *hsdM* deletion mutant, an overlap PCR using Phusion DNA polymerase was performed of the region upstream of *hsdM* with primers 93 and 108, and the region downstream of *hsdM* with primers 96 and 109. The resulting PCR products were used as template for a second PCR reaction with primers 96 and 93. The final product was ligated into pJRS233 [15]. The resulting plasmid was transformed into GAS strain 854 by electroporation and subjected to allelic replacement as previously described [16]. Genotype of the mutant strain 854Δ*hsdM* was confirmed by PCR amplification of the *hsdM* locus, and phenotype was verified by qRT-PCR for loss of *hsdM* expression.

**Table S1.**
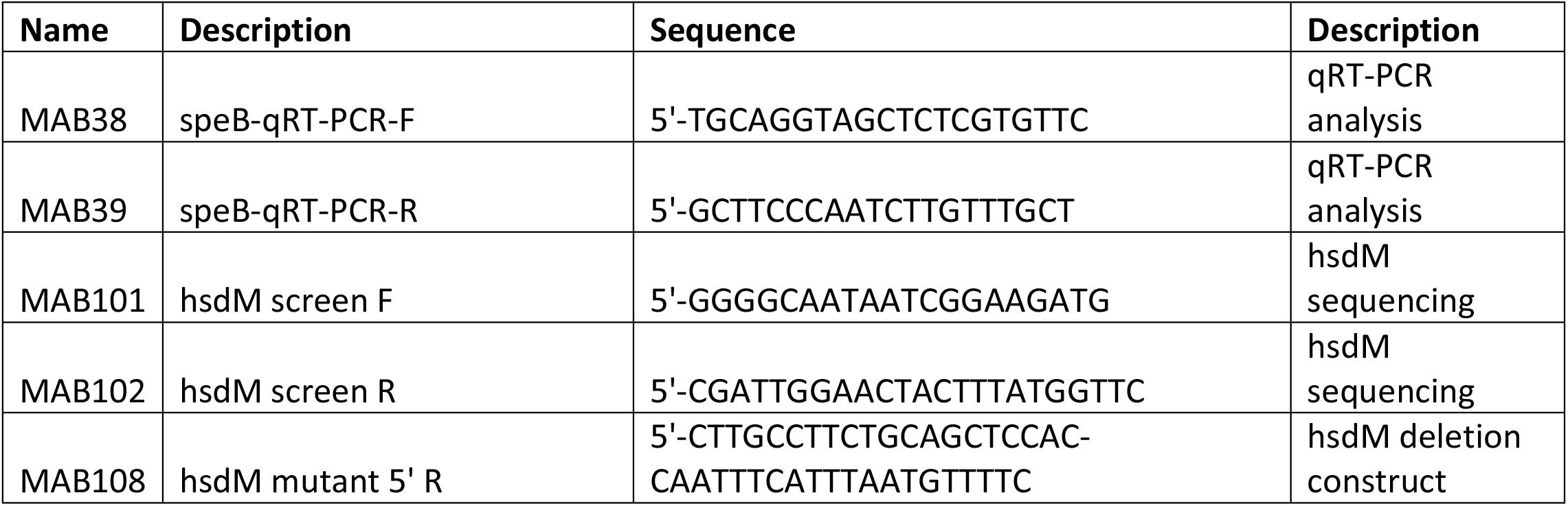
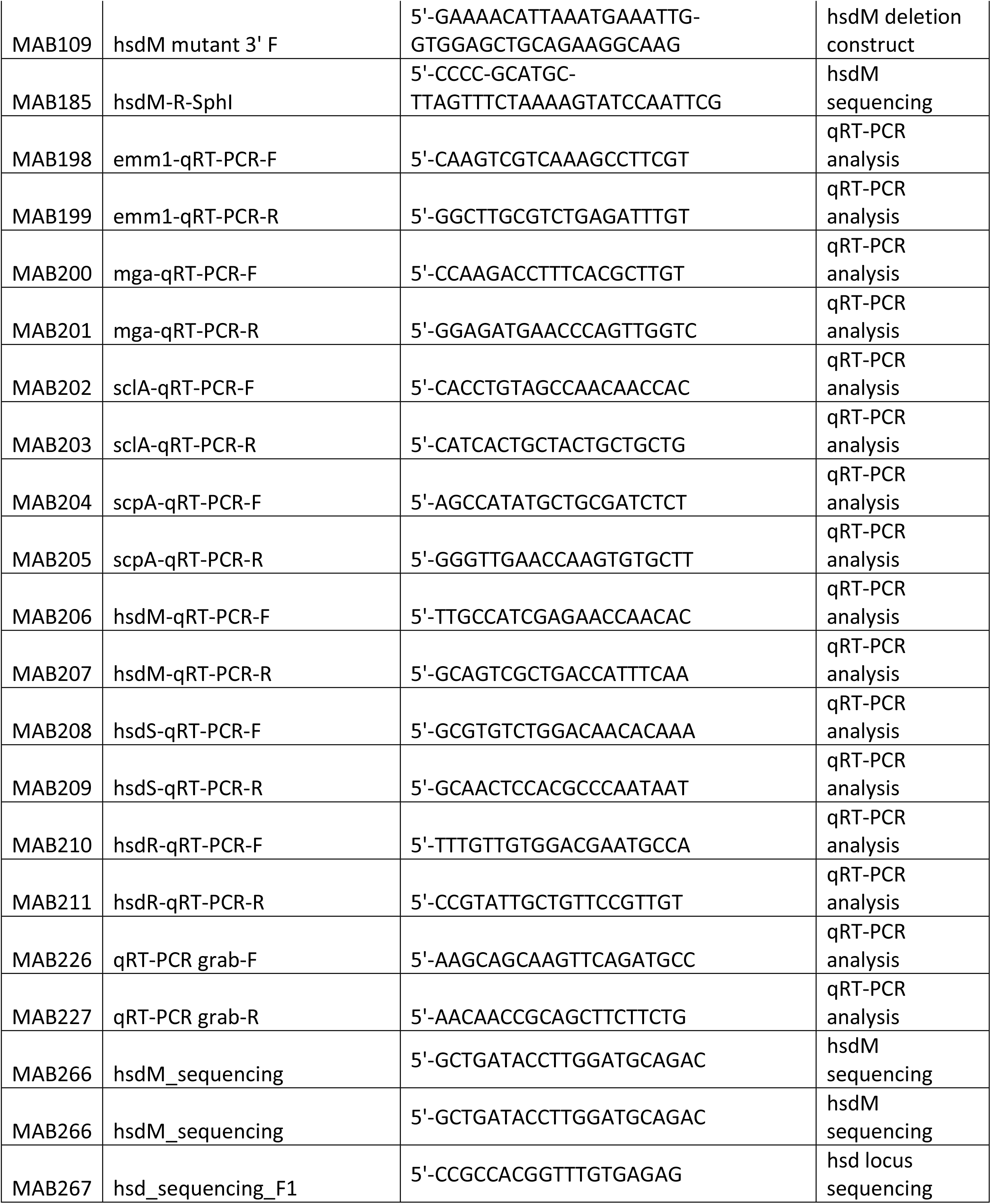
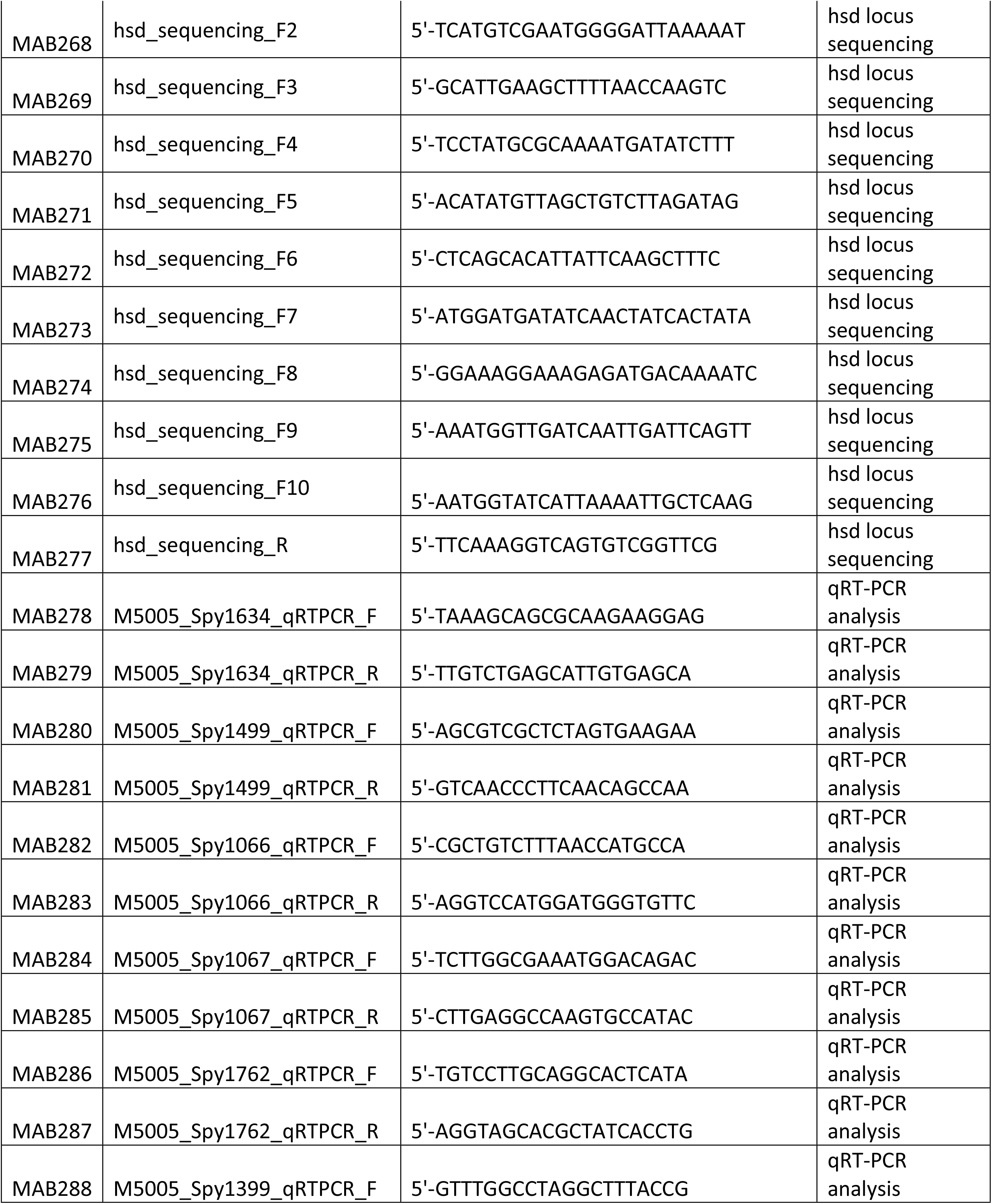
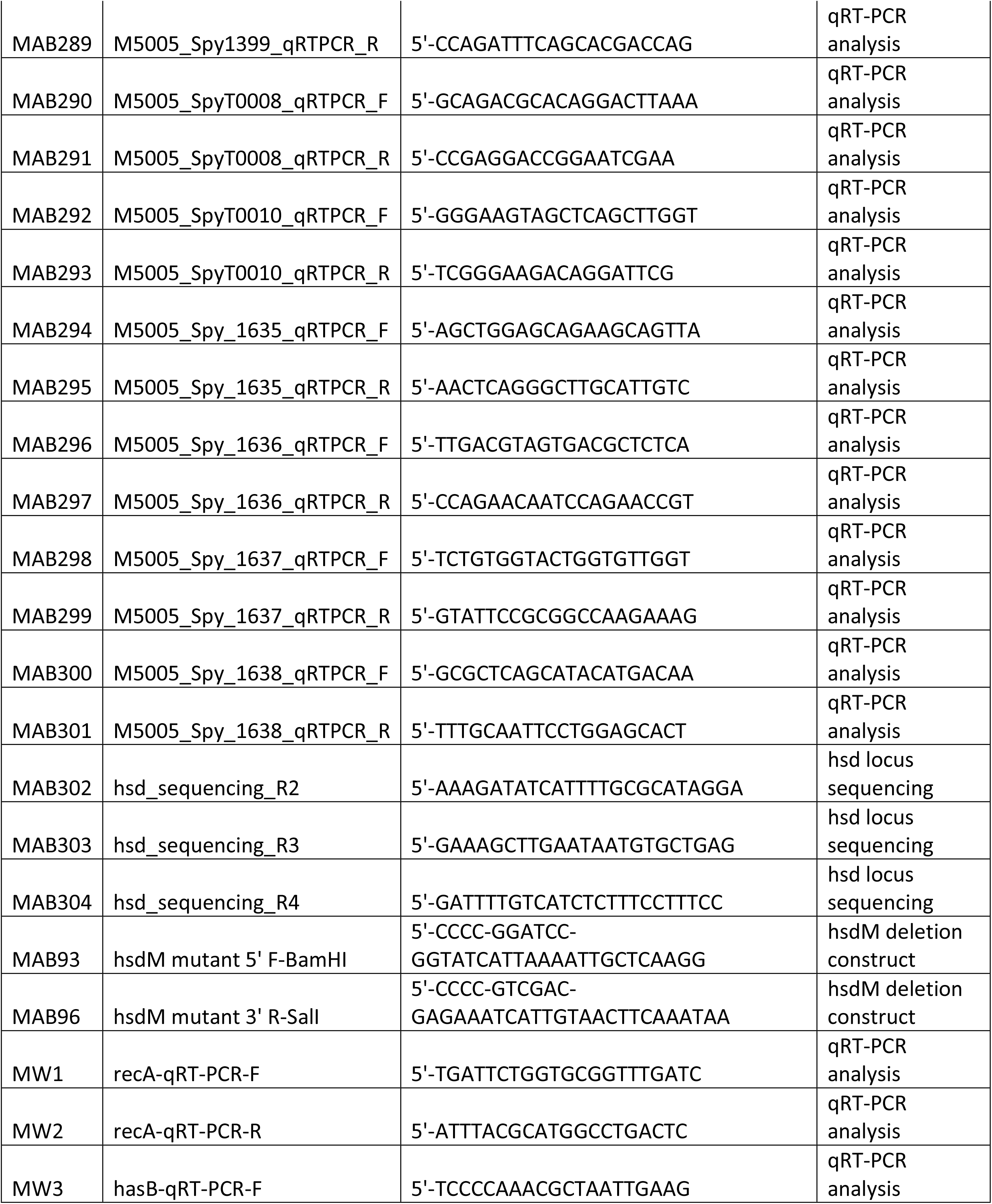
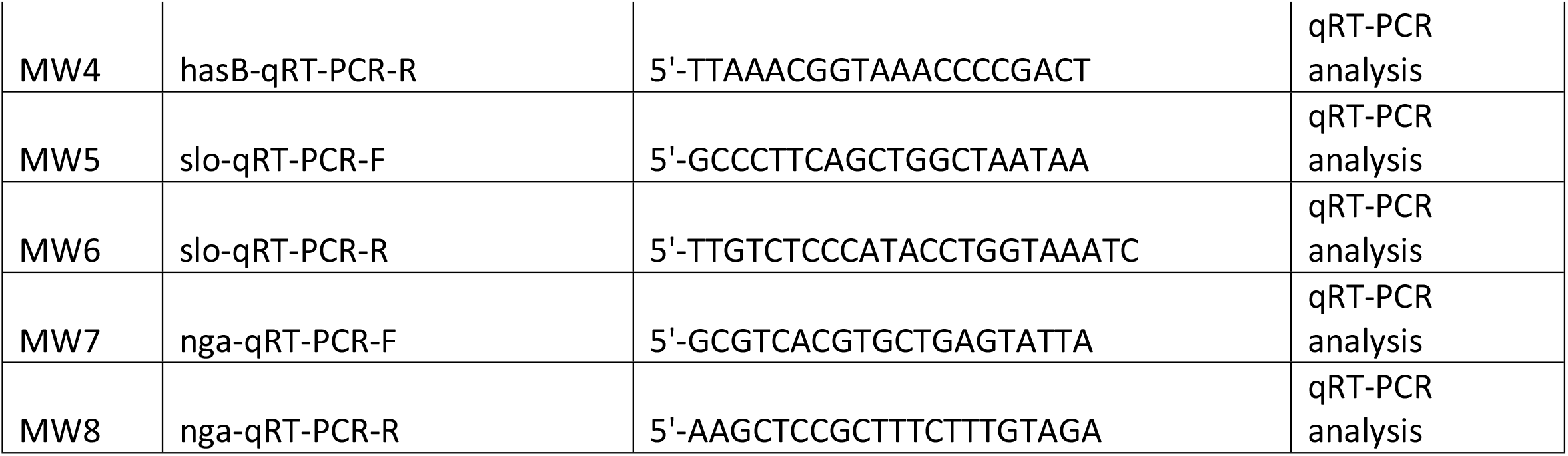
Oligonucleotide primers used in this study.

### RNA isolation and qRT-PCR

GAS were grown in THY broth, and cells were harvested at either mid-exponential (A_600_ 0.5), or early stationary (A_600_ 1) growth phase. Total RNA extraction from bacterial cells was performed using an RNeasy mini kit (Qiagen) according to the manufacturer’s instructions. RNA concentration and purity were determined using a NanoDrop spectrophotometer ND-1000 (Thermo Fisher Scientific). cDNA was generated using the High Capacity RNA-to-cDNA Kit (Applied Biosystems) according to the manufacturer’s instructions.

Quantitative RT-PCR was performed on a QuantStudio 3 System (Applied Biosystems) using the PowerUp SYBER Green Master Mix (Applied Biosystems). Primers are listed in Table S1. Triplicate assays were performed for each gene tested with 1 ng total template. Expression levels of target genes were normalized to *recA* (M5005_Spy_1799), and then further normalized to the wild type strain expression levels to obtain a final ΔΔCt value. These values were used to calculate relative expression changes for the mutant strain compared to the parental wild type strain. Data were reported as mean log_2_ fold-change.

### Transformations and quantification of transformation efficiency

Plasmid pDL278 was isolated from *E. coli* strain NEB5α (New England Biolabs) using a QIAfilter Plasmid Midi Prep (Qiagen). Wild type strain 854 or 854Δ*hsdM* was subjected to electroporation with 1 μg of plasmid DNA. After one hour of recovery, the reaction was serially diluted and spread on TH agar supplemented with 5% defibrinated sheep blood and 100 μg/ml spectinomycin. Colonies, representing transformants, were counted after overnight incubation at 37°C.

### RNA extraction for RNA-seq

GAS were grown in THY broth to mid-exponential phase (A_600_ 0.4) and collected by centrifugation. Cell pellets were resuspended in 0.5 mL Trizol reagent (ThermoFisher Scientific) and were transferred to 2 mL FastPrep tubes (MP Biomedicals) containing 0.1 mm Zirconia/Silica beads (BioSpec Products) and shaken for 90 seconds at 10 m/sec speed using the FastPrep-24 5G (MP Biomedicals). After addition of 200 µl chloroform, each sample tube was mixed thoroughly by inversion, incubated for 3 minutes at room temperature, and subjected to centrifugation for 15 minutes at 4°C. The aqueous phase was mixed with an equal volume of 100% ethanol, transferred to a Direct-zol spin plate (Zymo Research), and RNA was extracted according to the Direct-zol protocol (Zymo Research).

### Generation of RNA-seq data

Illumina cDNA libraries were generated using a modified version of the RNAtag-seq protocol [17] Briefly, 0.5 to 1 μg of total RNA was fragmented, depleted of genomic DNA, dephosphorylated, and ligated to DNA adapters carrying 5’-AN_8_-3’ barcodes of known sequence with a 5’ phosphate and a 3’ blocking group. Barcoded RNAs were pooled and depleted of rRNA using the RiboZero rRNA depletion kit (Epicentre). Pools of barcoded RNAs were converted to Illumina cDNA libraries in 2 main steps: (i) reverse transcription of the RNA using a primer designed to the constant region of the barcoded adaptor with addition of an adapter to the 3’ end of the cDNA by template switching using SMARTScribe (Clontech) as described [18] (ii) PCR amplification using primers whose 5’ ends target the constant regions of the 3’ or 5’ adaptors and whose 3’ ends contain the full Illumina P5 or P7 sequences. cDNA libraries were sequenced on the Illumina [Nextseq 2500] platform to generate paired-end reads.

### RNA-seq Data Analysis

Paired-end sequencing reads were mapped to the *S. pyogenes* MGAS5005 genome (NCBI RefSeq NC_007297) using bowtie2 version 2.3.1 [19]. Reads that mapped to annotated genes were counted using HTSeq version 0.10.0 and analysis of differential gene expression was conducted using DESeq2 version 1.20.0. Reported genes had a two-fold greater or lesser abundance than wild type, all with an adjusted *P*-value of 0.05 or less.

### Modeling of HsdM from GAS

Sequence of HsdM from MGAS5005 (accession number WP_010922657.1) was submitted to I-TASSER to be compared with the cryo-EM structure of HsdR_2_M_2_S_1_-Ocr^IS^ from EcoR124I (PBD 7BST) [20,21]. Molecular graphics and analyses were performed with UCSF Chimera, developed by the Resource for Biocomputing, Visualization, and Informatics at the University of California, San Francisco [22].

## Acknowledgements

We thank Jorge Velarde for assistance in modeling the GAS HsdM structure and generating the structural figure. RNA-seq libraries were constructed and sequenced at the Broad Institute of MIT and Harvard by the Microbial ‘Omics Core and Genomics Platform, respectively.

